# Regenerative Failure Following Rat Neonatal Chorda Tympani Transection is Associated with Geniculate Ganglion Cell Loss and Terminal Field Plasticity in the Nucleus of the Solitary Tract

**DOI:** 10.1101/503326

**Authors:** Louis J. Martin, Amy H. Lane, Kaeli K. Samson, Suzanne I. Sollars

**Affiliations:** Department of Psychology, University of Nebraska at Omaha, Omaha, NE, USA

**Keywords:** taste, gustatory, glossopharyngeal nerve, greater superficial petrosal nerve, nerve section, regeneration

## Abstract

Neural insult during development results in recovery outcomes that vary dependent upon the system under investigation. Nerve regeneration does not occur if the rat gustatory chorda tympani nerve is sectioned (CTX) during neonatal (≤ P10) development. It is unclear how chorda tympani soma and terminal fields are affected after neonatal CTX. The current study determined the impact of neonatal CTX on chorda tympani neurons and brainstem gustatory terminal fields. To assess terminal field volume in the nucleus of the solitary tract (NTS), rats received CTX at P5 or P10 followed by chorda tympani label, or glossopharyngeal (GL) and greater superficial petrosal (GSP) label as adults. In another group of animals, terminal field volumes and numbers of chorda tympani neurons in the geniculate ganglion (GG) were determined by labeling the chorda tympani with DiI at the time of CTX in neonatal (P5) and adult (P50) rats. There was a greater loss of chorda tympani neurons following P5 CTX compared to adult denervation. Chorda tympani terminal field volume was dramatically reduced 50 days after P5 or P10 CTX. Lack of nerve regeneration after neonatal CTX is not caused by ganglion cell death alone, as approximately 30% of chorda tympani neurons survived into adulthood. Although the total field volume of intact gustatory nerves was not altered, the GSP volume and GSP-GL overlap increased in the dorsal NTS after CTX at P5, but not P10, demonstrating age-dependent plasticity. Our findings indicate that the developing gustatory system is highly plastic and simultaneously vulnerable to injury.

**Highlights:**

- The volume of chorda tympani central processes is severely diminished after the nerve is sectioned in 5- or 10-day-old rats.
- Compared to adult injury, neonatal denervation produced greater loss of terminal field and ganglion cell bodies.
- About 30% of chorda tympani neurons survive early section, suggesting lack of regeneration is not caused by cell loss alone.
- The organization of intact gustatory terminal fields is altered by early chorda tympani nerve section.

## Introduction

Outcomes following nervous system injury vary greatly depending on the animal’s age during insult (Kolb et al., 1987; Taylor and Alden, 1997; Kempler et al., 1999; Crowe et al., 2012; Semple et al., 2013). Compared to adults, young animals have a higher degree of plasticity following neural injury (Gómez-Pinilla et al., 1986; Burgess and Villablanca, 1986; Villablanca et al., 1986; Himes and Tessler, 1989; Johansson and Arvidsson, 1994; Renehan et al., 1994; Rhoades et al., 1997), but this is not always associated with better recovery (Giza and Prins, 2006; Anderson et al., 2011). In the peripheral nervous system, nerve injuries are often more disruptive in developing, compared to mature, animals (Aldskogius and Arvidsson, 1978; Aldskogius et al., 1985; Chiaia et al., 1987; Hosley et al., 1987; Sollars, 2005; Omelian et al., 2016).

The taste system is an excellent model to study developmental differences in plasticity and injury response. Injury to the chorda tympani (CT), the nerve that innervates taste buds in fungiform papillae and anterior foliate papillae, leads to strikingly different outcomes depending on the animal’s age. Adult rats receiving chorda tympani transection (CTX) experience a loss of taste buds on the anterior tongue and deficits in taste-guided behavior that are largely reversed following nerve regeneration (St. John et al., 1995; Kopka et al., 2000). When CTX occurs in rats at postnatal day 10 (P10; neonatal) or younger, however, the CT does not regenerate (Sollars and Bernstein, 2000). Long-term changes to the taste system occur, including loss of almost all anterior tongue taste buds (Sollars, 2005), development of abnormal taste preferences (Sollars and Bernstein, 1996), and altered activity to NaCl stimulation in intact gustatory nerves (Martin and Sollars, 2015). It is unclear why the CT does not regenerate, but there may be large-scale death of CT neurons in the geniculate ganglion (GG). Approximately 20% of CT neurons are lost after adult CTX (Schuler et al., 2004), and ganglion cell death following denervation is much more severe for neonatal rats than adults in the dorsal root (Himes and Tessler, 1989) and trigeminal ganglia (Aldskogius and Arvidsson, 1978; Chiaia et al., 1987).

Terminal fields of taste nerves in the nucleus of the solitary tract (NTS) display a high degree of plasticity. The CT, glossopharyngeal (GL; the nerve that innervates taste buds in circumvallate and posterior foliate papillae), and greater superficial petrosal (GSP; which innervates taste buds in the soft palate, nasoincisor duct, and geschmacksstreifen) terminal fields normally prune in volume by roughly one-third between P15 and adulthood (Sollars et al., 2006; Mangold and Hill, 2008). The size of gustatory terminal fields is influenced by diet (Pittman and Contreras, 2002; May and Hill, 2006; Sollars et al., 2006; Thomas and Hill, 2008), taste receptor protein expression (Skyberg et al., 2017; Sun et al., 2017), and neurotrophic factors (Sun et al., 2015; 2018).

There is evidence that afferents compete for terminal space during development. Early postnatal whisker lesions cause degeneration of innervating neurons (Waite and Cragg, 1979) and uninjured nerve fibers expand into the denervated area of the brainstem (Johansson and Arvidsson, 1994; Renehan et al., 1994). In the gustatory system, transection of the GL and GSP causes the CT terminal field to increase in size fourfold, thereby mimicking its neonatal state (Corson and Hill, 2011). Interestingly, this phenomenon is not developmentally-dependent, as transection at P15, P35, or adulthood produces the same effects. This age-independent plasticity is surprising given the large differences in gustatory plasticity across age following CTX (Akaike et al., 1965; Markison et al., 1995; Sollars and Bernstein, 1996; Sollars, 2005; Martin and Sollars, 2015).

Minor loss of CT terminal field and CT neurons occurs after adult CTX (Shuler et al., 2004; Reddaway et al., 2012), but it is not clear how the CT itself is impacted by neonatal CTX. The present study was designed to determine the effects of neonatal CTX on CT neurons and intact nerve terminal fields of the GL and GSP. To examine this, gustatory nerve terminal fields were labeled following neonatal CTX, and CT neurons in the GG were examined after neonatal or adult CT denervation. We found that neonatal CTX caused a profound loss of CT terminal field and ganglion cells and altered the organization of intact gustatory terminal fields.

## Experimental Procedures

### Subjects

Female Sprague-Dawley rats (n = 47) were used for these experiments. Rats were bred in the University of Nebraska at Omaha vivarium. The day of birth was designated P0, and animals were weaned at P25. Rats were socially housed in clear Plexiglas cages where they had free access to food pellets (Teklad or Labdiet) and tap water. Animals were kept on a 12:12 light-dark cycle. All procedures were approved by the local Institutional Animal Care and Use Committee.

### Chorda Tympani Transection

Rats received surgery at P5, P10, or P50 as previously described (Sollars, 2005). Rats were anesthetized with Brevital^®^ (methohexital sodium; 60 mg/kg, i.p.) and the right CT was accessed through an incision in the ventromedial neck. To perform CTX, the right CT was either cut with a pair of fine surgical scissors or crushed proximal to its juncture with the lingual nerve and evulsed, removing approximately 5 mm of the nerve. The CT was cut using scissors only in animals receiving nerve label (DiI) at the time of CTX (see below). Evulsion was used for all other CTX surgeries. The impact of P10 CTX on fungiform papillae number and morphology is the same regardless of the nerve is cut or evulsed (Sollars et al., 2002), so no differences in regeneration would be expected. Neonatal CT evulsion does not alter the number of lingual nerve fibers (Sollars et al., 2002), indicating that the lingual nerve is not impacted by CTX. In Sham animals, the CT was visualized, but left intact. The wound was then closed with a suture, and rats were placed on a heating pad. Upon recovery from the anesthetic, animals were returned to their home cage.

### Chorda Tympani Nerve Label

#### Biotinylated Dextran Amine

Rats received CT label 50-51 days after P5 CTX (n =5), P10 CTX (n=5), or Sham surgery at P5 (n = 3) or P10 (n = 2). Animals were anesthetized with a mixture of ketamine (73 mg/kg) and xylazine (7 mg/kg) given i.p., and the right tympanic bulla was accessed ventrally using blunt forceps. The CT was visualized in the tympanic bulla and cut at the point where it crosses the ossicles. The proximal end of the CT was carefully moved onto a shelf of bone and a small piece of cotton soaked in dimethyl sulfoxide (DMSO) was placed on the nerve for 15 s. Crystals of 3 kDa biotinylated dextran amine (BDA; Invitrogen, BDA-3000) were liberally applied onto the cut CT, and left for at least 15 min. The tympanic bulla was then filled with Kwik-Sil (World Precision Instruments) to keep the CT and dye in place. The wound was closed with 2-3 sutures. Animals were placed on a heating pad during recovery from the anesthetic. Once sternal recumbency was regained, animals were returned to their home cages.

#### DiI

Because it has been suggested that BDA transport to the brain becomes impaired after long-term neural injury (Reddaway et al., 2012) the long-lasting tracer DiI (1,1’-Dioctadecyl-3,3,3’,3’-Tetramethylindocarbocyanine Perchlorate; D3911, Invitrogen) was applied in another group of animals at the time of CTX at P5 or P50 (n = 6/group). Since there is no long-term loss of CT volume after adult CTX (Reddaway et al., 2012), the rats labeled at P50 served as controls. Animals were anesthetized with Brevital^®^ (50 mg/kg, i.p.) and the CT was accessed in the same manner as the nerve transection procedure. The nerve was cut with fine surgical scissors and DMSO was applied to the central stump of the CT with a small piece of cotton. Crystals of DiI were placed onto the nerve, and the rat was maintained in a supine position for at least 15 min. Animals recovered from the anesthetic and were returned to their home cage.

### Glossopharyngeal and Greater Superficial Petrosal Nerve Label

GL and GSP terminal fields were simultaneously labeled in rats 50 days after P5 CTX (n = 5) or P10 CTX (n = 5) or in P55 – P60 intact controls (n = 5). Following anesthesia with Brevital^®^ (50 mg/kg, i.p.), a hole was made in the ventral aspect of the right tympanic bulla. The tensor tympani muscle was removed, and a portion of the overlying temporal bone was removed to expose the GSP. DMSO and crystals of BDA were applied to the cut nerve in the same manner as CT nerve label. At least 15 min after BDA application, the tympanic bulla was filled with Kwik-Sil. The GL was then located just ventral and medial to the tympanic bulla. After cutting the GL and applying DMSO, crystals of 3 kDa tetramethylrhodimine dextran (TMR; Invitrogen, D3308) were placed onto the central stump of the nerve. The rat was kept in place for at least 15 min to ensure adequate uptake of the neural tracer. The wound was sutured closed and animals were placed on a heating pad until recovery from the anesthetic.

### Tissue Processing

Rats were sacrificed either two days following label with dextrans (BDA or TMR) or 50-51 days following DiI label. Animals were overdosed with a mixture of ketamine/xylazine and transcardially perfused with a modified KREBS solution (pH = 7.3) followed by 4% paraformaldehyde dissolved in an acetate buffer and 4% paraformaldehyde in a borate buffer (for double-labeled animals) or 8% paraformaldehyde (all other rats). Brains were extracted and stored in 8% paraformaldehyde overnight. Horizontal sections (50 μm) were obtained using a vibratome filled with 0.1 M phosphate buffered saline (PBS). The horizontal axis was chosen because the full extent of the gustatory NTS can be observed in relatively few sections. This type of sectioning also allowed us to distinguish the dorsal NTS, the area where experimentally-induced changes in gustatory terminal field size are often most extreme (Lasiter and Kachele, 1990; King and Hill, 1991; Krimm and Hill, 1997; Pittmann and Contreras, 2002; Mangold and Hill, 2007; Mangold and Hill, 2008; Sun et al., 2015).

For double-labeled tissue, free-floating sections were placed in a solution containing Streptavidin Alexa Fluor 488 (Molecular Probes; 1:500 in PBS/0.1% triton mixture) for 2 hr to visualize BDA. TMR did not require further processing. Tissue was then rinsed (2 x 5 min) in PBS before placing on gelatin-coated slides. After air-drying, slides were coverslipped with DPX.

For CT terminal field labeled with BDA, free-floating sections were processed for diaminobenzidine (DAB) visualization (Corson et al., 2012). Sections were rinsed in 0.01 M PBS (pH: 7.4, 5 x 5 min) before being placed in a 4% avidin-biotin complex solution (Vector Laboratories) in 0.01 M PBS with 0.1% triton at 4°C overnight. Following another 0.01 M PBS rinse (1 x 10 min and 2 x 15 min), sections were placed in a 0.01 M PBS solution containing 0.05% DAB, 0.125% nickel ammonium sulfate, and 0.0015% H_2_O_2_ for 10-20 min. Sections were washed in distilled water (3 x 1 min and 2 x 5 min), mounted on gelatin-coated slides, and coverslipped with DPX after air-drying.

### Terminal Field Tracing

Terminal field volumes were assessed by tracing the area containing bouton-bearing fibers using Neurolucida (MBF Bioscience). This technique allowed us to identify the space in the NTS occupied by terminal field while excluding solitary tract axons. Tracings were performed by a researcher blind to experimental condition. Terminal fields were traced with a 10X (x 1.6) objective for all brains except those in which the CT was labeled with BDA. Given the small amount of CT terminal field often evident after neonatal CTX, these CT tracings were performed using a 20X (x 1.6) objective. DAB-stained CT terminal fields were visualized with brightfield illumination, whereas GL and GSP terminal fields and DiI-labeled CT terminal fields were viewed using epifluorescence. The area of each traced section was multiplied by the section thickness, 50 µm, to obtain terminal field volume. Total terminal fields volumes and the area of overlap between GL and GSP terminal fields were assessed.

The NTS was divided into three contiguous regions along the dorsal-ventral axis, similar to previous research (Pittmann and Contreras, 2002; May and Hill, 2006; Sollars et al., 2006; Mangold and Hill, 2007; 2008). The intermediate zone of the NTS was identified first by finding sections in which tracts oriented in the rostral-caudal plane passed through the gustatory terminal fields in the rostral NTS. Visualization of these tracts was often enhanced by dark field settings. Three to four sections were identified as intermediate for each animal. The dorsal and ventral NTS zones were classified as sections that were dorsal or ventral to the intermediate zone, respectively. Given the anatomical orientation of the NTS, dorsal sections are most caudal and ventral sections are more rostral. See Figure 1 for an example of the NTS zones. For brains in which the CT was labeled with DiI, the terminal field was not divided into zones because labeled field appeared in relatively few sections.

**Figure 1.**
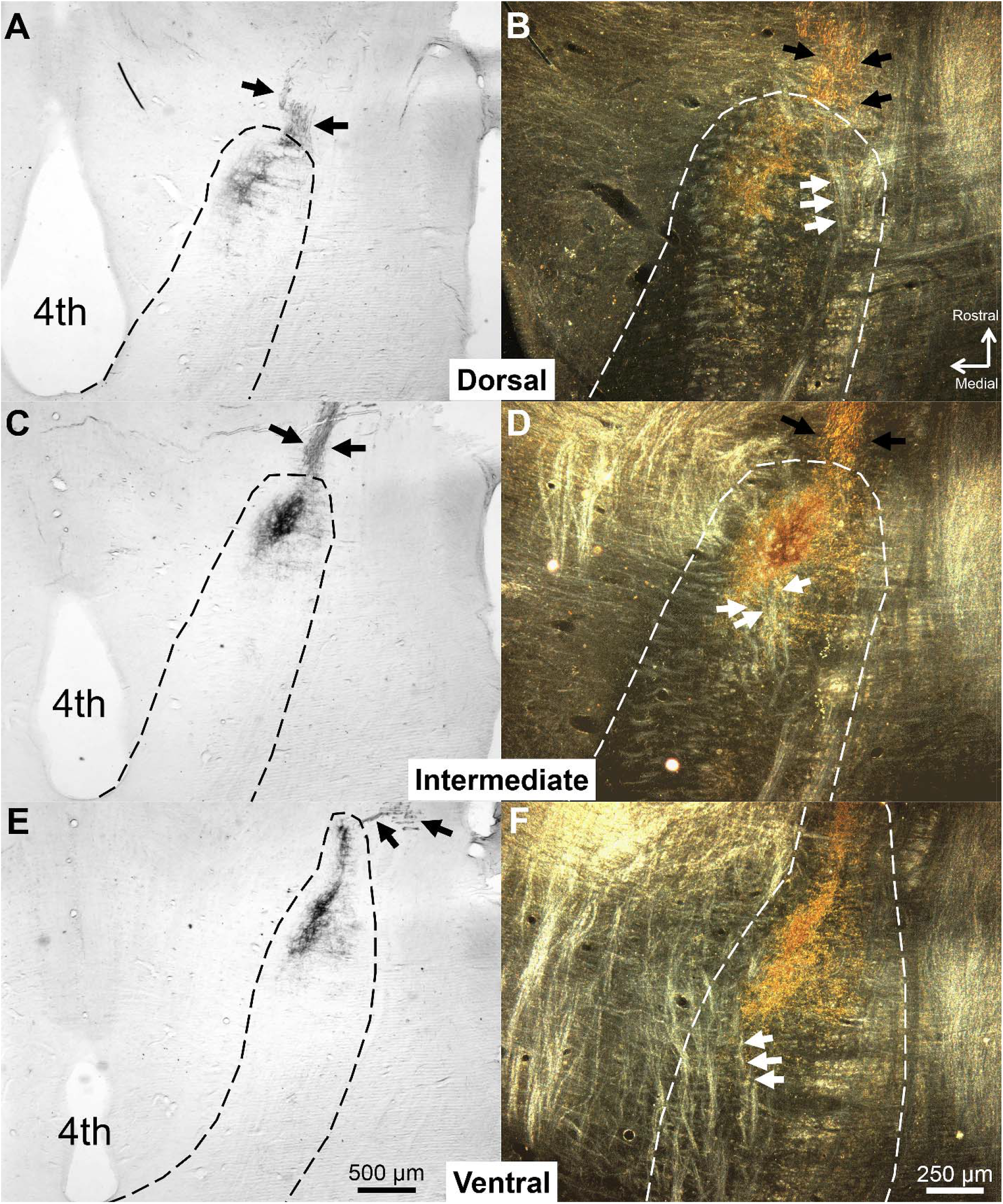
CT terminal field in the dorsal, intermediate, and ventral NTS in a Sham rat labeled with BDA at P60. The NTS was sectioned in the horizontal plane. (A, C, E) Low-magnification, brightfield images of the NTS (dotted line). (B, D, F) Higher magnification images of the NTS are shown using darkfield illumination. In the darkfield images, arrows point to tracts near the CT terminal field that are lighter than surrounding tissue. These tracts are oriented in the rostral-caudal plane and are lateral to gustatory terminal fields in the dorsal NTS. As the NTS progresses ventrally, these tracts situate more medially and move through gustatory terminal fields. We identified sections in which these tracts were within the terminal field area as the intermediate NTS. Black arrows indicate chorda tympani axons projecting toward the NTS. 4^th^ = Fourth ventricle

### Geniculate Ganglion

To determine the extent of cell loss after neonatal CTX, CT neuron counts in the GG were performed in a portion of the animals receiving CT label with DiI at P5 (n = 5) or P50 (n = 4). After perfusion, the right GG was extracted with the facial nerve and a large portion of the GSP attached. GG were cryoprotected using 30% sucrose and flash-frozen in −80° C methylbutane. Tissue was cut into 10 µm sections with a cryostat and mounted onto gelatin-coated slides. DiI-labeled GG cells were visualized with an epifluorescent microscope. Cells containing DiI and the entire ganglion were traced using Neurolucida before staining the tissue with 1% cresyl violet and coverslipping. Using Neurolucida, the tracings from the DiI label were overlaid onto the stained tissue and cells were tracked across sections and counted by a researcher blind to experimental condition. This technique yielded the number of CT cells that survived neonatal or adult CTX.

To assess the long-term effects of neonatal CTX on cell survival, an additional group of 5 rats received unilateral CTX at P10. Rats were perfused and both the right and left GG were extracted between 567 and 937 (*M* = 643) days after surgery. Central immune responses in neonatal rats do not differ between the NTS contralateral to CTX and the NTS of unoperated or Sham-sectioned control rats (Riquier & Sollars, 2017). Therefore, neurons in the GG contralateral to CTX are unlikely to be damaged, making the contralateral GG an appropriate control. Ganglia were cryosectioned at 10 µm, stained with 1% cresyl violet, and coverslipped. Because these rats did not receive DiI label at the time of transection and nerve labels transport poorly in the gustatory nerves of aged rats (S.I. Sollars, unpublished observations), total ganglion cell counts were assessed. Since each neuron appeared on multiple sections, a soma was counted only when the nucleolus was clearly visible (Konigsmark, 1970; Miller et al., 1978; Srinivasan & Stevens, 2018). Cell counts were compared between the right (CTX) and left (intact) ganglia. See Table 1 for a summary of the experimental design.

**Table 1.** Experimental Design

| Experiment | Nerve Label | Surgical Conditions | Time of Nerve Label | Tissue Extraction | Variable Assessed |
| --- | --- | --- | --- | --- | --- |
| CT Terminal Field | BDA | P5 CTX (n = 5)<br>P10 CTX (n = 5)<br>Sham (n = 5) | 50-51 Post Surgery | 48 hr After Label | Terminal Field Volume<br>Dorsal-Ventral Extent of Field |
|  | Dil | P5 CTX (n = 6)<br>P50 CTX (n = 6) | Immediately After CTX | 50-51 Days Post-Surgery | Terminal Field Volume |
| CT Ganglion Cell Counts | Dil | P5 CTX (n = 5)<sup>a</sup><br>P50 CTX (n = 4)<sup>a</sup> | Immediately After CTX | 50-51 Days Post-Surgery | Number of Labeled GG Neurons |
| GL-GSP Nerve Label | GL – TMR<br>GSP – BDA | P5 CTX (n = 5)<br>P10 CTX (n = 5)<br>No surgery (n = 5) | 50 Days After Surgery | 48 hr After Label | Terminal Volume<br>Terminal Field Overlap |
| GG Neuron Counts (aged rats) | N/A | P10 CTX (n = 5) | N/A | 567-937 Days Post-Surgery | GG Neurons Ipsilateral to CTX<br>vs. GG Neurons Contralateral to CTX |

a. These animals are a subset of those used to assess CT terminal field using Dil.

### Verification of Surgery

To ensure that the CT was transected in CTX rats, the dorsal surface of the tongue was stained with 0.1% methylene blue and fungiform papillae were inspected under a microscope. Following CTX at P10 or younger, most fungiform papillae are no longer detectable, and the majority of the few remaining papillae have altered morphology (Sollars and Bernstein, 2000; Sollars, 2005). Identifying the morphology and number of fungiform papillae is, thus, an easy measure of the success of surgery. Surgical success was verified in all cases.

### Data Analysis

The effect of CTX on CT terminal field volume for BDA-labeled animals was determined using a two-way repeated-measures ANOVA: surgery (P5 CTX, P10 CTX, Sham) x NTS zone (dorsal, intermediate, ventral). A One-Way ANOVA compared the size of the CT terminal field in the dorsal-ventral plane (number of sections with BDA-labeled CT terminal field x section thickness) between groups. The impact of CTX on GL and GSP terminal fields was tested with a three-way repeated-measures ANOVA: Nerve (GL or GSP) x Surgery (P5 CTX, P10 CTX, intact) x NTS Zone (dorsal, intermediate, ventral). The effect of surgery on GL/GSP overlap was determined with a two-way repeated-measures ANOVA: Surgery (P5 CTX, P10 CTX, intact) x NTS Zone (dorsal, intermediate, ventral). Bonferroni post-hoc tests were used to compare the effect of surgical condition on total terminal field volume as well as the volume within each NTS zone for each nerve. Differences in total GG neuron counts in aged rats were assessed with a paired *t*-test.

For all comparisons, each surgical group was tested for normality. The number of DiI-labeled ganglion cells and the volume of DiI-labeled terminal field were not normality distributed within surgical groups (P5 label vs. P50 label). Therefore, we made these comparisons with non-parametric Mann-Whitney U tests. For all other tests, the data from either the majority or all of the surgical groups were normally distributed. The alpha level for each test was .05.

## Results

### Chorda Tympani Terminal Field after Chorda Tympani Transection

CT terminal field volumes assessed using BDA as the neural tracer were substantially diminished following neonatal CTX (Figs 2, 3). There was a significant main effect of surgery (*F*(2,12) = 36.485, *p* < .001) on CT terminal field volume. Overall CT volume and volumes within all NTS zones were significantly lower for transected animals compared to controls (each *p*< .01). There were no significant differences in total CT volume or volume within each zone between P5 CTX and P10 CTX rats (each *p* > .10). There was a non-significant trend for an interaction between surgery and NTS zone on CT volume (*F*(4,24) = 2.706, *p* = .054). Although CT volumes appear most diminished in the dorsal zone following transection, the amount of terminal field was low throughout the NTS for animals sectioned at P5 or P10. The extent of CT terminal field in the dorsal-ventral plane was not significantly different among surgical groups (*F*(2,12) = 2.235, *p* = .150).

**Figure 2.**
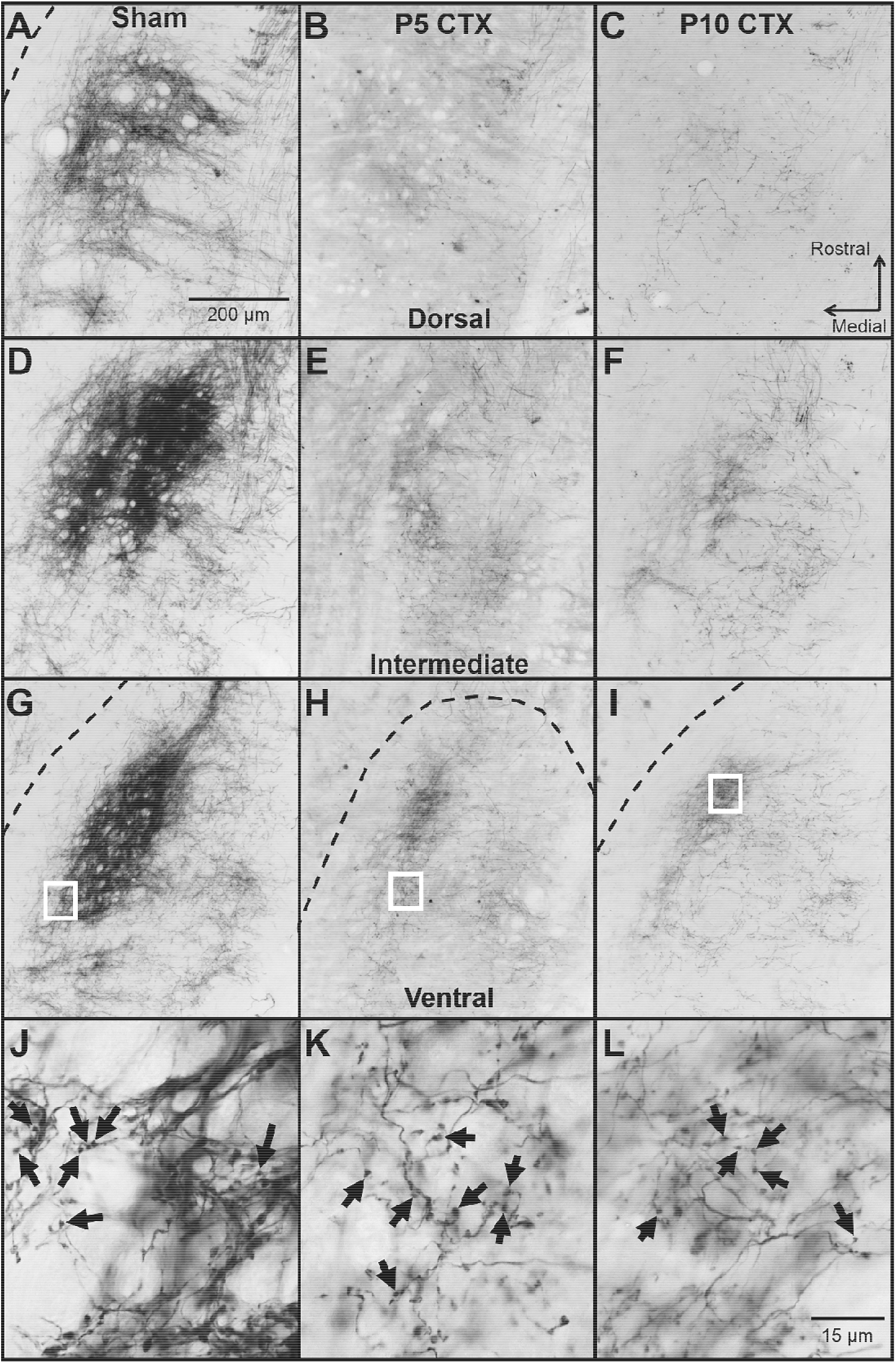
Representative images of CT terminal field in the dorsal (A-C), intermediate (D-F), and ventral (G-I) NTS. Images are from an animal receiving Sham surgery at P10 (A, D, G, J), CTX at P5 (B, E, H, K), or CTX at P10 (C, F, I, L). The CT was labeled with BDA 50-51 days after surgery. The borders of the NTS are indicated with a dashed line in cases when the entire image is not within the NTS. In transected animals, spared CT fibers were observed in regions typically occupied by the control CT, even in the outer borders of the NTS. J-L shows a high magnification view of the CT terminal field in the ventral NTS. These images were taken from the areas outlined by white boxes in G-I. Examples of labeled boutons are shown with black arrows.

**Figure 3.**
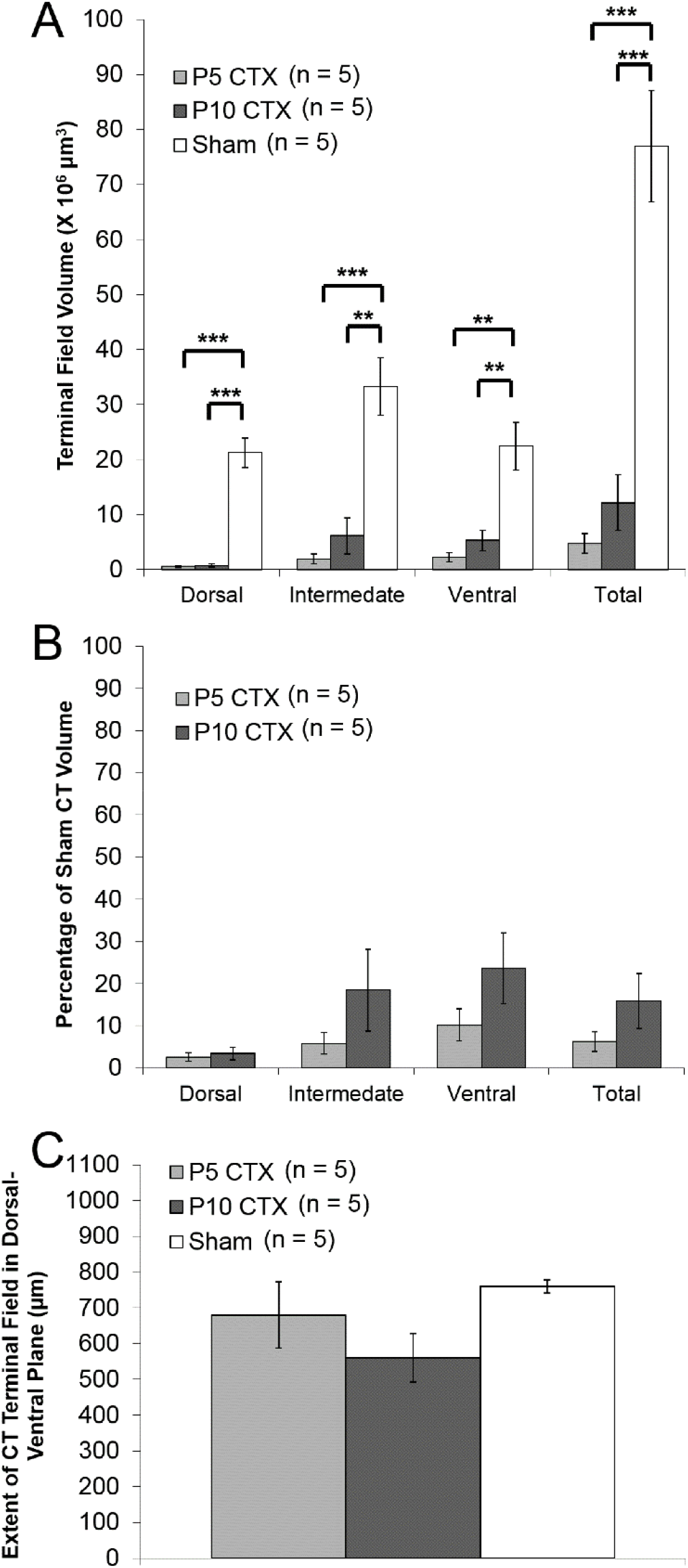
CT terminal field data for adult rats labeled with BDA following neonatal surgery. (A) Average (±SEM) CT terminal field volumes for animals that received CTX at P5, CTX at P10, or Sham surgery. (B) Percent of Sham CT volume for P5 CTX and P10 CTX rats (C) Size of CT terminal field in the dorsal-ventral plane. ** *p* < .01 *** *p* < .001

One potential explanation for reduced CT volume after neonatal surgery is that BDA does not transport properly when applied to a nerve that was previously transected (Reddaway et al., 2012). To address this possibility, the CT was labeled with DiI at the time CTX occurred. In all animals, DiI staining was found in the geniculate ganglion and salivatory cells of the medulla, indicating proper transport. Terminal field staining was less widespread than with BDA, but the densest portion of the CT terminal field was evident (Fig 4). Animals labeled at P5 had significantly lower CT volumes than rats labeled at P50 (*U* = 35, *p* = .004). This result is consistent with our findings using BDA and suggest that CT terminal fields are severely diminished following neonatal CTX.

**Figure 4.**
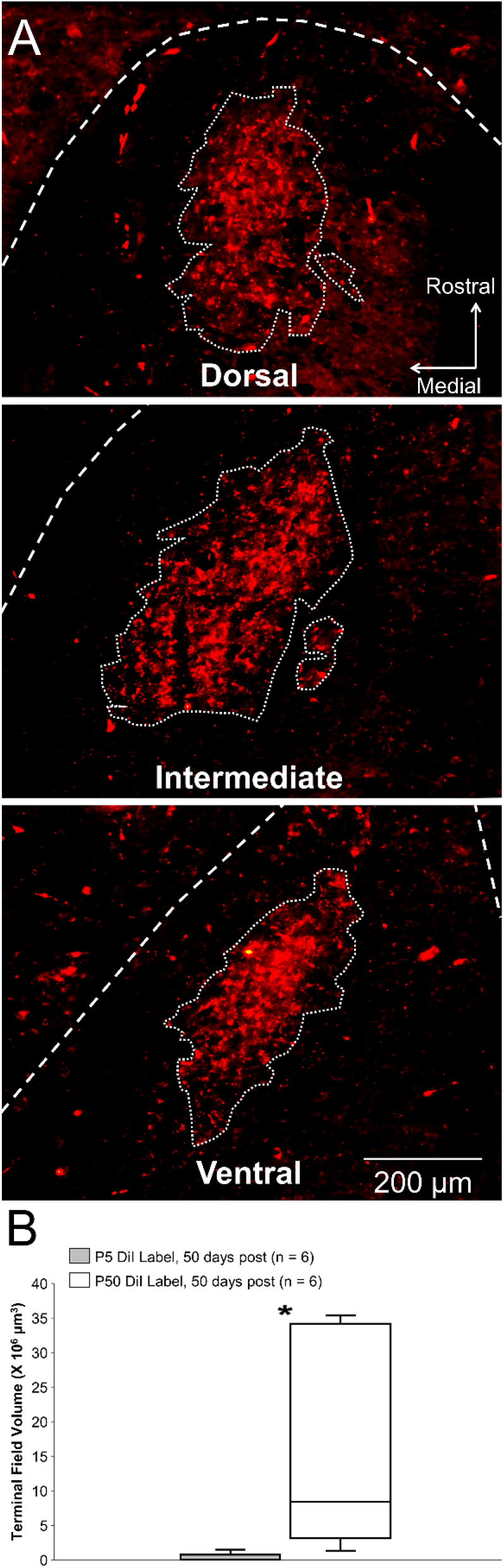
(A) Example images of CT terminal field in the NTS after DiI label at P50. Dashed lines indicate the terminal field border. Unlike staining with BDA, fine CT fibers were not observed with DiI. (B) Box-and-whisker plot displaying CT terminal field volume in animals labeled with DiI at P5 or P50. * *p* < .01

### Glossopharyngeal and Greater Superficial Petrosal Terminal Field After Chorda Tympani Transection

Early CT denervation altered the organization of intact gustatory terminal fields (Fig 5). There was no significant main effect of surgery on terminal field volume (*F*(2,24) = 0.523, *p* = .599) and no significant surgery by nerve interaction (*F*(2,24) = 0.181, *p* = .836). These findings suggest that early CTX does not impact the total terminal field volume of either the GL or the GSP. However, there was a significant interaction between surgery and NTS zone (*F*(4,48) = 6.492, *p* < .001). Post-hoc tests revealed that in the dorsal zone, the volume of the GSP terminal field was significantly higher for P5 CTX rats compared to intact rats (*p* = .039). There was a tendency for GSP volumes in the dorsal zone to be higher after P5 CTX compared to P10 CTX although this difference was not significant (*p* =.072). In the ventral zone, rats receiving CTX at P10 had significantly lower GSP volumes than intact rats (*p* = .040). For the GL, there were non-significant trends for higher volumes after P5 CTX compared to P10 CTX (*p* = .052) and intact rats (*p* = .067).

**Figure 5.**
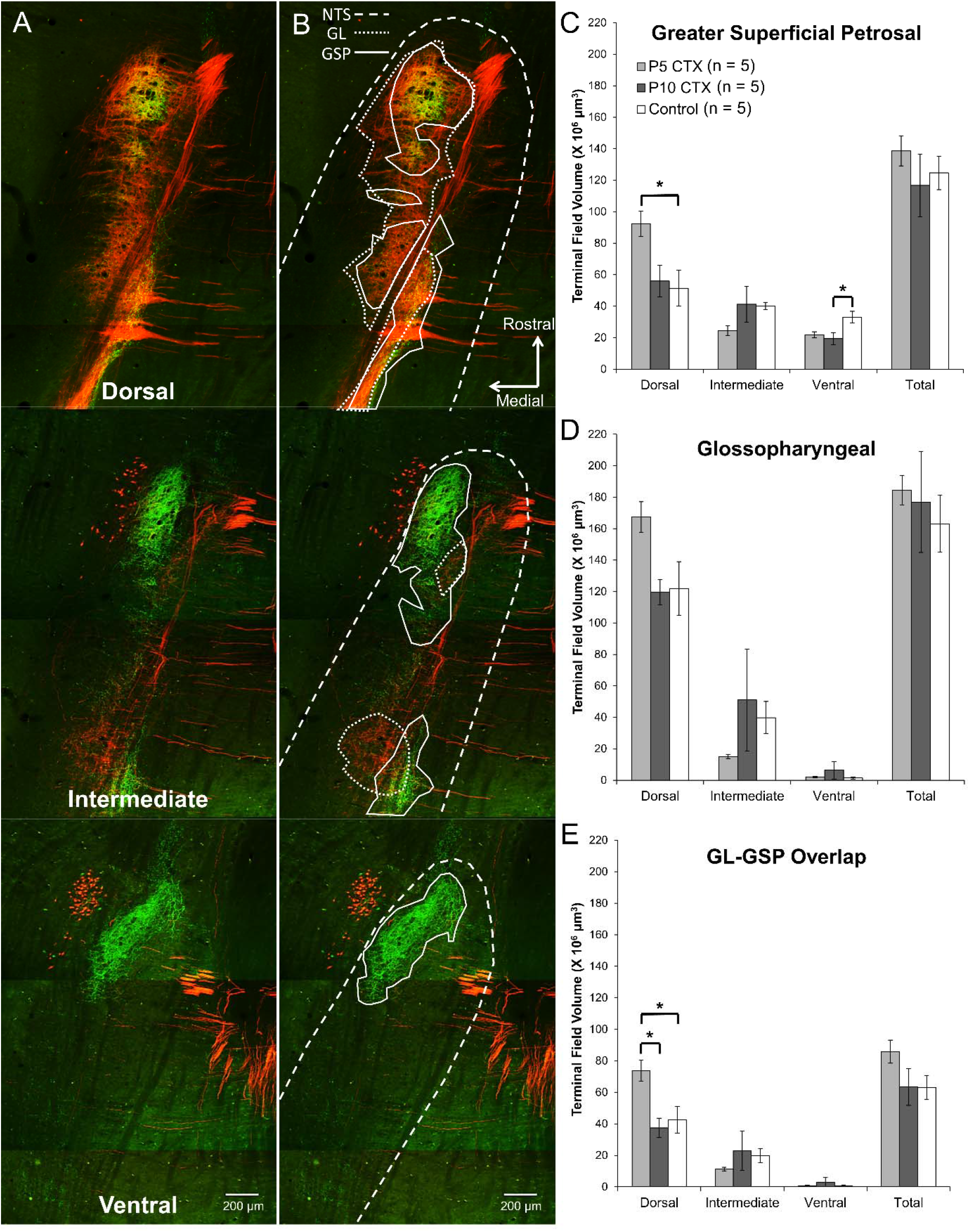
(A) Images of GL (red: TMR-labeled) and GSP (green: BDA-labeled) terminal fields in a double-labeled rat. (B) The same images as (A) with the NTS borders drawn with a dashed line, the GL terminal field outlined with a dotted line, and the GSP terminal field designated with a solid line. (C) Mean (±SEM) terminal field volume for the GSP in adult rats receiving CTX at P5, CTX at P10, or no surgery. (D) Average (±SEM) GL terminal field volume (E) Mean (±SEM) overlap between the GL and GSP terminal fields * *p* < .05

**Figure 6.**
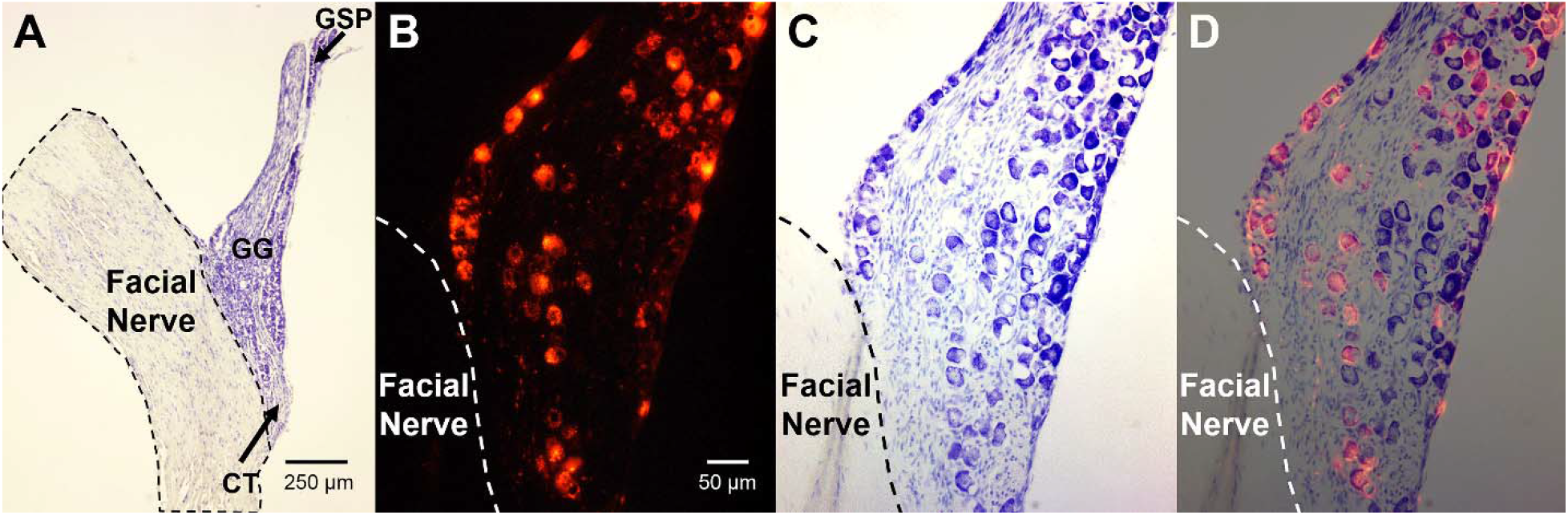
Images of the geniculate ganglion (GG). (A) Example creysl-violet-stained GG from an adult (P57) rat that received CTX at P5. Nerves were not labeled, and this animal was not used for data analysis. The facial nerve is outlined with a dashed line. Black arrows indicate the area from which the CT and GSP enter the GG. (B) GG from a different rat than (A) that received DiI label at P50 and was sacrificed 50 days later. The location of the facial nerve is indicated by the dashed line. (C) The same tissue as (B) stained with cresyl violet. (D) Merged image of (B) and (C).

There was no main effect of surgery on the volume of overlap between the GL and GSP terminal fields (*F*(2,12) = 2.036, *p* = .173), but there was a significant interaction between surgery and NTS zone (*F*(4,24) = 4.267, *p* = .009). In the dorsal zone, the volume of overlap was significantly higher for P5 CTX rats compared to controls (*p* = .030) and P10 CTX animals (*p* = .012). There were no differences between surgical groups in the intermediate or ventral NTS (each *p* > .10). CTX appears to alter the typical pattern of overlap between the GL and GSP, but only if the CT is denervated very early postnatally, as P10 CTX was not effective. The fact that P5 CTX was more effective at altering terminal field organization than P10 CTX suggests that the differences seen here are not a general effect of surgery, but specific to the timing of CT denervation.

Rats that received DiI label at P5 had significantly fewer stained GG cells compared to animals labeled at P50 (Table 2; *U* = 20, *p* = .016). Of those DiI-labeled animals for which both terminal field and GG cell count data were available (P5, n = 4; P50, n = 4) there was a significant positive correlation between the number of labeled ganglion cells and CT terminal field volume (*r* = .856, *p* = .007). In aged rats receiving unilateral CTX at P10, total CV cell counts were significantly lower in the ganglion ipsilateral to CTX compared to the ganglion contralateral to CTX (Table 3; *t*(4) = 10.75, *p* < .001). The ipsilateral ganglion had an average of 396 fewer neurons than the intact, contralateral GG. Because the GG contains approximately 500 CT neurons (Miller et al., 1978; Lasiter and Kachele, 1990; Shuler et al., 2004), one would expect about 100 CT cells in these rats. These findings suggest that CT cell death is greater when denervation occurs in early development, but many CT neurons survive transection.

**Table 2.** DiI-labeled CT neurons 50-51 Days After Surgery

| P5 CTX | P50 CTX |
| --- | --- |
| 248 | 479 |
| 121 | 260 |
| 116 | 297 |
| 135 | 318 |
| 106 | - |
| **145.2 (26.1)** | **338.5 (48.3)** |

Mean number of cells shown in bold and standard error of the mean in parentheses

**Table 3.**
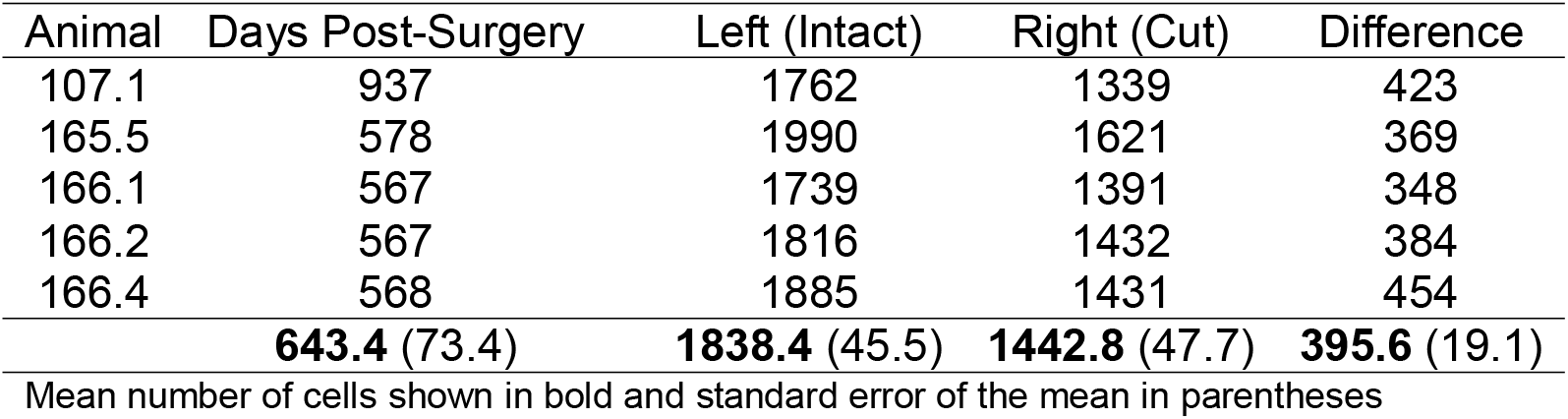
Total GG Cells after P10 CTX

## Discussion

In the present study, we found that neonatal CTX causes a severe loss in CT terminal field volume and a greater reduction of CT cells in the GG compared to adult nerve section. The lack of regeneration after neonatal denervation is not caused by complete CT neuron death, as approximately 30% of CT neurons survive P5 CTX. Although neonatal CTX did not change the total volume of the GL or GSP terminal field, loss of the CT altered the pattern of organization. There was an increased GSP volume and higher GSP-GL overlap in the dorsal zone after CTX at P5, but not P10. These findings indicate that the effects of CTX on intact terminal fields depends on the timing of loss, with younger rats exhibiting greater plasticity.

Compared to control animals and rats sectioned as adults (Reddaway et al., 2012), neonatal CTX caused a substantial reduction in CT terminal field. Despite the absence of CT regeneration, many central nerve fibers were preserved. Neonatal CTX did not impact the extent of the CT terminal field in the dorsal-ventral plane, and bouton-bearing fibers were evident in all areas of the NTS typically seen in adults – including a small amount of labeling in the caudal NTS. A similar preservation of central projections is observed in nociceptive neurons when innervation of the skin is prevented (Olson and Luo, 2018). Thus, a permanent removal of the distal CT may not alter the development and maintenance of spared central fibers.

Consistent with findings from the somatosensory system (Aldskogium and Arvidsson, 1978; Chiaia et al., 1987; Himes and Tessler, 1989), we found that ganglion cell loss is greater for neonates than adults following transection. We observed an average of 145 CT cells in adult rats that received P5 CTX. We also found that after P10 CTX, there were 396 fewer neurons in the GG ipsilateral to CTX compared to the contralateral GG in adult animals. The lack of CT regeneration after neonatal CTX is associated with a large, but not exceptional, amount of neuronal loss. Because the GG contains approximately 500 CT neurons (Miller et al., 1978; Lasiter and Kachele, 1990; Shuler et al., 2004), about 100-150 neurons appear to survive neonatal CTX – even 1.8 years after denervation. Although the degree of cell death is high, it is comparable to loss following P0 transection of the infraorbital nerve or sciatic nerve, where 40% and 25% of neurons survive, respectively (Chiaia et al., 1987; Himes and Tessler, 1989). Both these nerves regenerate following neonatal denervation (Chiaia et al., 1987; Schmalbruch, 1987), suggesting that the absence of CT regeneration is not simply the result of inadequate neuronal survival. Preventing regeneration in the infraorbital nerve following P0 transection does not influence cell loss (Chiaia et al., 1987). Thus, regenerative failure following neonatal CTX may not contribute to cell death.

Given the preservation of roughly 30% of CT neurons, regenerative failure may to be a result of inadequate targets that facilitate regeneration. Global knockout of brain-derived neurotrophic factor (BDNF) in adult mice prevents regeneration of the CT without affecting ganglion cell death (Meng et al., 2017). Following CTX in adult mice, BDNF expression increases in preserved taste buds and is retained in tissue adjacent to the taste bud (Meng et al., 2017), likely providing attractant signals for regenerating CT fibers. Although a portion of taste buds and most fungiform papillae are spared after CTX in adult rats (Segerstad et al., 1989; Li et al., 2015), almost all taste buds and ~2/3rds of fungiform papillae are lost following neonatal CTX (Sollars and Bernstein, 2000; Sollars et al., 2002; Sollars, 2005). The majority (80%) of remaining fungiform papillae are “filiform-like,” highly keratinized structures that likely cannot grow new taste buds. These changes to taste buds and papillae on the anterior tongue may reduce the presence of chemoattractants such as BDNF, impeding regeneration. To test whether the preservation of papillae is sufficient to facilitate CT regeneration, one could examine taste buds in the anterior foliate papillae following neonatal CTX. The CT innervates a portion of anterior foliate taste buds (Miller et al., 1978) and degeneration of the foliate papillae would not be expected following neonatal CTX. Although we have not been able to observe the CT outside the tympanic bulla after neonatal CTX (Sollars and Bernstein, 2000), a small number of fibers may regenerate and support foliate taste buds.

The number of CT neurons that survive neonatal CTX is higher than we anticipated given the minimal terminal field remaining. Reddaway and colleagues (2012) found a 63% reduction in CT terminal field volume after adult CTX when using BDA as a tracer, but not when DiI was applied at the time of CTX. This finding suggests that long-term neural injury may impair the transport of BDA. To our knowledge, it is currently not possible to selectively stain all CT fibers without a nerve label. Nevertheless, we found long projections of CT fibers in the NTS with numerous boutons extending to the limits of the typical field size, suggesting successful BDA tract tracing after P5 and P10 CTX. The sparsity of fibers, not the intensity of BDA label, was most notable in our study, supporting our finding that neonatal CTX causes a large decrease in CT field size that is not an artifact of the method of tract tracing. The CT terminal field size we observed using BDA after neonatal CTX is much lower than that seen after adult CTX (Reddaway et al., 2012) and we found smaller DiI-labeled terminal field after P5 CTX compared to CT denervation at P50. Additionally, the CT in the tympanic bulla of neonatally transected animals was noticeably much smaller than that of Sham rats at the time of BDA label.

Neonatal CTX altered the organization, but not the total size of GL and GSP terminal fields. CTX at P5 caused greater changes to intact terminal fields compared to denervation at P10, even though there was no significant difference in terminal field degeneration between animals sectioned at P5 and P10. These findings suggest that the malleability of GL and GSP terminal fields decreases across development. In contrast, when the GL and GSP are transected in P15, P25, or adult rats, the CT expands fourfold regardless of age at transection (Corson & Hill, 2011). While not precisely comparable, since ages examined in the current study are younger than those examined by Corson and Hill (2011), terminal field plasticity across development appears to vary among gustatory nerves. Experimental manipulations that influence gustatory nerve function or terminal field size are most effective in influencing the CT, followed by the GSP and then the GL (Hill and Przekop, 1988; King and Hill, 1991; Sollars and Hill, 2000; May and Hill, 2006; Sun et al., 2015; Skyberg et al., 2017; Sun et al., 2018). These findings are consistent with the results of the present study and suggest that the CT is especially plastic as compared to other gustatory nerves.

In summary, CTX at P5 or P10 causes substantial long-term reductions in CT neurons in the GG and in the CT terminal field when compared to sham control and adult rats. The high degree of CT neuron death is comparable to the neonatal somatosensory system where regeneration of the nerve occurs (Chiaia et al., 1987; Schmalbruch, 1987; Himes and Tessler, 1989). Thus, regenerative failure of the CT after neonatal CTX is likely due to factors other than loss of neurons. Following CTX at P5, but not P10, the increase in GSP volume and GSP-GL overlap in the dorsal NTS indicates that terminal field plasticity decreases across neonatal development in these nerves. Taken together with our earlier findings of extreme vulnerability of neonatal fungiform taste buds and papillae to CT transection, and the synergistic interactions between the CT and lingual nerve that are particularly pronounced during neonatal ages (Sollars, 2005; Omelian et al, 2016), the present results further demonstrate the unique characteristics of the developing gustatory system.

## Acknowledgements

Funding: this work was supported by the National Institutes of Health [DC04846] and the University of Nebraska at Omaha Office of Research and Creative Activities. These institutions did not influence the design of the study, collection of data, or interpretation of results. The authors thank Andrew Riquier and Kristi Apa for comments on an earlier version of this manuscript and Jonathan Santo for his assistance with data analysis.

## Author Contributions

Designed study: LM, AL, KS, SS

Performed experiments: LM, AL, KS, SS

Analyzed data: LM, SS

Wrote paper: LM, SS

## Glossary

BDA: biotinylated dextran amine
CT: chorda tympani nerve
CTX: chorda tympani transection
DAB: diaminobenzidine
DiI: 1,1’-Dioctadecyl-3,3,3’,3’-Tetramethylindocarbocyanine Perchlorate
DMSO: dimethyl sulfoxide
GL: glossopharyngeal nerve
GG: geniculate ganglion
NTS: nucleus of the solitary tract
P: postnatal day
PBS: phosphate-buffered saline
TMR: tetramethylrhodamine dextran

